# Estrogen receptor beta in the central amygdala regulates the deleterious behavioral and neuronal consequences of repeated social stress in female rats

**DOI:** 10.1101/2022.10.03.509933

**Authors:** Cora E. Smiley, Brittany S. Pate, Samantha J. Bouknight, Megan J. Francis, Alexandria V. Nowicki, Evelynn N. Harrington, Susan K. Wood

## Abstract

While over 95% of the population has reported experiencing extreme stress or trauma, females of reproductive age develop stress-induced neuropsychiatric disorders at twice the rate of males. This suggests that estrogen may facilitate neural processes that increase stress susceptibility and underlie the heightened rates of these disorders, like depression and anxiety, that result from stress exposure in females. However, there is contradicting evidence in the literature regarding estrogen’s role in stress-related behavioral outcomes. Estrogen signaling through estrogen receptor beta (ERβ) has been traditionally thought of as anxiolytic, but recent studies suggest estrogen exhibits distinct effects in the context of stress. Furthermore, ERβ is found abundantly in many stress-sensitive brain loci, including the central amygdala (CeA), in which transcription of the vital stress hormone, corticotropin releasing factor (CRF), can be regulated by an estrogen response element. Therefore, these experiments sought to identify the role of CeA ERβ activity during stress on behavioral outcomes in naturally cycling, adult, female Sprague-Dawley rats. Rats were exposed to an ethological model of vicarious social stress, witness stress (WS), in which they experienced the sensory and psychological aspects of an aggressive social defeat encounter between two males. Following stress, rats exhibited stress-induced anxiety-like behaviors in the marble burying task, and, finally, brain analysis revealed increased ERβ and CRF specifically within the CeA following exposure to stress cues. Subsequent experiments were designed to target this receptor in the CeA using microinjections of the ERβ antagonist, PHTPP, prior to each stress session. Sucrose preference, acoustic startle, and marble burying tasks determined that blocking ERβ in the CeA during WS prevented the development of depressive-,anxiety-like, and hypervigilant behaviors. Additionally, brain analysis revealed a long-term decrease of intra-CeA CRF expression in PHTPP-treated WS rats compared to vehicle. These experiments indicate that ERβ signaling in the CeA, through its effects on CRF, contribute to the development of negative valence behaviors that result from exposure to repeated social stress in female rats.

## 1. Introduction

Social stressors are one of the most commonly experienced and highly impactful forms of stress in the clinical population. Exposure to stress is a risk factor for the development of many neuropsychiatric and physical health disorders, including depression, anxiety, and cardiovascular disease (Pêgo, Sousa, Almeida, & Sousa, 2010). As a leading cause of disability, such stress-related conditions pose a significant burden on both the patient and society. While the overall incidence of mental health disorders is alarming, affecting around 30% of the population, females exhibit over twice the rate of neuropsychiatric conditions, including anxiety and depression when compared to males, possibly due to increased exposure and vulnerability to stress (Sandanger, Nygård, Sørensen, & Moum, 2004). Importantly, this incidence is not correlated with differences in demographic factors such as marital status, employment, or number of children (Klose & Jacobi, 2004; Steel et al., 2014). Therefore, neurobiological factors inherent to females may underly this increased prevalence of mental health disorders, especially following stress exposure.

Despite a large body of literature suggesting that, in healthy patients, estrogen mediates protective effects under non-stressful conditions, recent advances in the field have established estrogen as a factor that may promote susceptibility to the detrimental consequences of stress (J. E. Finnell et al., 2018a; Hokenson et al., 2021; Morgan & Pfaff, 2001; Morgan, Schulkin, & Pfaff, 2004). Higher levels of circulating estrogen have been shown to increase physiological and neuroendocrine responses to stress (T. D. Lund, Hinds, & Handa, 2006), and in post-menopausal women without circulating estrogen, injections of estrogen lead to a heightened response to laboratory administered social stressors (Newhouse et al., 2010). Further, elevated estrogen in rodent models, as a consequence of microbead neuronal implants or cycle stage, promotes an exaggerated release of stress hormones, specifically a critical regulator of the stress response, corticotropin releasing factor (CRF), and amplifies neuronal activation of brain regions responsible for the production of these hormones (Babb, Masini, Day, & Campeau, 2013; Handa, Burgess, Kerr, & O’Keefe, 1994; T. D. Lund et al., 2006; Viau, Bingham, Davis, Lee, & Wong, 2005; Zuloaga, Heck, De Guzman, & Handa, 2020). Thus, estrogen may promote this susceptibility through its direct actions on the neuroendocrine stress response (Heck & Handa, 2019; Zuloaga et al., 2020).

Estrogen has also been shown to directly increase the expression of CRF mRNA and heighten behavioral responses to stressful stimuli (A. M. Jasnow, J. Schulkin, & D. W. Pfaff, 2006). Importantly, the CRF gene contains an estrogen response element which allows nuclear estrogen receptor (ER) activation to modulate CRF transcription (Ni & Nicholson, 2006; Vamvakopoulos & Chrousos, 1993). Using a novel model of vicarious social defeat stress, referred to here as witness stress (WS), we previously reported a stress-evoked increase in CeA CRF among intact but not ovariectomized female rats (J. E. Finnell et al., 2018a). Furthermore, this sensitized CRF response specific to intact females was accompanied by increased negative valence behavioral responses. Along with evidence indicating that estrogen-mediated effects on CRF represent a promising mechanism for increased stress susceptibility among females, these findings highlight the need to determine the specific neural mechanisms underlying this phenomenon.

Due to these known interactions with CRF, estrogen signaling within specific stress-sensitive brain regions may be instrumental in exacerbating stress sensitivity in females. The central amygdala (CeA) is one such region that contains a prominent CRF network that is activated in response to stress (Herringa, Nanda, Hsu, Roseboom, & Kalin, 2004; Kalin, Shelton, & Davidson, 2004; Pomrenze et al., 2019). This region is also critically involved in regulating behaviors related to hypervigilance, including the elevated plus maze (Tye et al., 2011) and acoustic startle response (Liang et al., 1992), as well as long-term negative responses to stress cues (Weera, Shackett, Kramer, Middleton, & Gilpin, 2021). Further, the CeA extends dense CRF projections to other stress-sensitive regions like the locus coeruleus (LC), which is known to be highly involved in integrating these signals to enhance the stress response (Howard, Carr, Hill, Valentino, & Lucki, 2008; Valentino & Van Bockstaele, 2008). Importantly, estrogen is also able to modulate cellular activity of the CeA by signaling through ER-beta (ERβ), the predominant ER subtype in this region (Osterlund, Kuiper, Gustafsson, & Hurd, 1998; Shughrue & Merchenthaler, 2001). While ERβ has traditionally been shown to be anxiolytic (Trent D. Lund, Rovis, Chung, & Handa, 2005), this effect is only exhibited in behavioral tasks among naïve rats without a history of stress (Le Moëne, Stavarache, Ogawa, S, & Ågmo, 2019). The dichotomous role of ERβ activity is revealed in rats that are exposed to stress prior to testing on measures of anxiety-like behaviors, suggesting that ERβ may, instead, have anxiogenic effects in the context of stress (Aaron M. Jasnow, Jay Schulkin, & Donald W. Pfaff, 2006; Lynch et al., 2014). Therefore, these experiments sought to identify a specific role for intra-CeA ERβ in negative behavioral responses to social stress that have been observed in female rats. Previous experiments in our lab have determined that cycling ovarian hormones are required for female susceptibility to social stress (J. E. Finnell et al., 2018b), thus these new studies were designed to explore the neuronal mechanisms responsible for estrogen-mediated behavioral responses. Specifically, these studies determined 1) how ERβ expression is altered in the CeA following social stress exposure in females, and 2) if pharmacological blockade of ERβ during stress prevents the development of maladaptive behavioral responses to repeated social stress.

## 2. Materials and Methods

### 2.1 Experimental Design

Two main experiments were completed under the hypothesis that, during stress exposure, estrogen signals through ERβ to promote CRF expression, leading to heightened stress responsivity in females (**Figure S1C**). Rats in Experiment #1 (**Figure 1A**, n=6/treatment group) were tested on a baseline marble burying task to establish pre-stress levels of anxiety-like behaviors prior to exposure to five days of witness stress (WS) or control handling (CON). Subsequently, marble burying tests were completed post-stress to determine the impact of social stress on this index of anxiety-like behavior, and, finally, tissue collection occurred following exposure to the WS/CON context (i.e., the same environment and cues originally experienced during WS/CON exposure). Rats in Experiment #2 (**Figure 1B**, n=12-15/treatment group) were implanted with indwelling bilateral microinjection cannulas into the CeA and received injections of the selective ERβ antagonist, PHTPP (10 μM in 10% DMSO in saline), or vehicle (10% DMSO in saline) one hour prior to each of the five daily WS/CON exposures (15 minutes/day for 5 consecutive days). Rats underwent pre- and/or post-stress behavioral assays, as described in the timeline in **Figure 1B**, including sucrose preference, acoustic startle, marble burying, and elevated plus maze, followed by tissue collection at rest.

**Figure 1.**
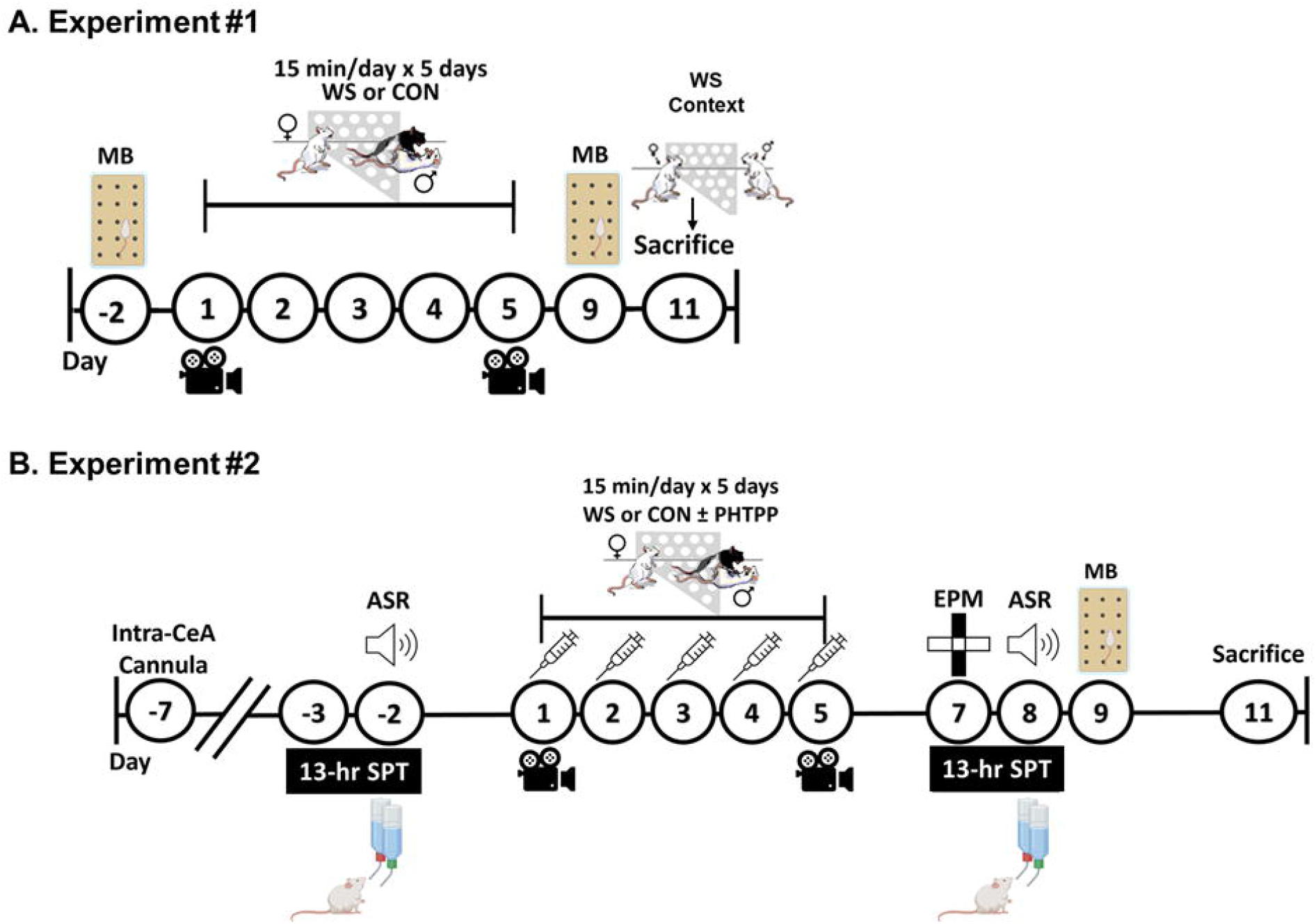
Experimental Timelines. Two separate experiments were completed to determine the role of CeA ERβ signaling in negative valence behavioral responses to repeated social stress exhibited by females. **A)** Rats in Experiment #1 were first assessed in a marble burying test (MB) to establish baseline levels of anxiety-like behavior, then were exposed to five consecutive days of WS/CON (15 minutes/day). Behaviors were assessed on days #1 (D1) and #5 (D5). WS/CON rats underwent a post-WS/CON MB test to determine the effects of WS compared to baseline measures of anxiety-like behavior. Finally, six days after the final WS/CON exposure, rats were re-exposed to the context in which they originally experienced WS (but in the absence of the resident) or CON for 15 minutes. Brains were collected 30 minutes after the start of WS/CON context exposure. **B)** For Experiment #2, all rats were implanted with indwelling bilateral cannulas in the CeA. Following surgical recovery, a pre-stress testing regimen commenced to assess baseline sucrose preference (SPT) and acoustic startle response (ASR). Subsequently, animals were exposed to five consecutive days of WS/CON (15 minutes/day) with local microinjection of either vehicle or the ERβ antagonist, PHTPP, one hour prior to each WS/CON session. Post-WS/CON behavioral testing consisted of the elevated plus maze (EPM), SPT, ASR, and MB, each separated by 24 hours, to determine if ERβ blockade (PHTPP) inhibited the development of associated negative valence behaviors. Tissue was collected at rest two days following the final behavioral test. CeA: Central Amygdala; ERβ: estrogen receptor beta

### 2.2 Animals

Female Sprague-Dawley rats (8-9 weeks and ∼150 g on arrival, witnesses and controls; Charles River, Raleigh, NC), male Sprague-Dawley rats (∼250 g on arrival, intruders; Charles River, Raleigh, NC), and male Long-Evans retired breeders (600-800 g, residents; Envigo, Dublin, VA) were housed individually in standard polycarbonate cages and maintained on a 12:12 light:dark cycle with lights on at 0700 hours with *ad libitum* access to standard rat chow and water, except during behavioral testing. Additionally, female rats that experienced control conditions were housed in a separate room from stressed female and all male rats. All studies were approved by the University of South Carolina’s Institutional Animal Care and Use Committee and maintained adherence to the National Research Council’s Guide for the Care and Use of Laboratory Animals.

### 2.3 CeA Cannulation Surgery

Rats in Experiment #2 were implanted with indwelling bilateral cannulas (26 ga., P1 Technologies, Roanoke, VA) aimed to terminate directly dorsal to the CeA (in mm from bregma and the surface of the dura, A/P: -2.3; M/L: ± 4.1, D/V: -6.5). Rats received post-operative analgesic injections (Flunazine, 0.25 mg/kg, s.c.) on the day of and day following surgery, along with nutritional supplementation (Bacon Softies, Bio-Serv), and underwent at least 7 days of recovery prior to the start of pre-stress testing.

### 2.4 Witness Stress/Control Handling

Females from both Experiment #1 and Experiment #2 were exposed to 15 minutes of WS/CON each day for five consecutive days. During each WS session, the female witness was placed behind a clear, perforated, acrylic partition in a protected region (10 × 5 cm) of a resident’s home cage. Immediately after placement of the witness, a male intruder was placed into the same side of the cage with the resident and social defeat commenced. This paradigm allows the witness to observe auditory, olfactory, and visual cues without physical aspects of social defeat (J. E. Finnell et al., 2018a). Matched witness-intruder pairs for all five WS sessions were exposed to a novel resident each day to prevent habituation. Prior to experimentation, residents were screened for appropriate levels of aggression. Residents that did not respond to an intruder or caused excessive physical injury to intruders were excluded. Controls were briefly handled then returned to their home cage, since we have previously shown that the presence of an intruder in the absence of social defeat produces no stress effect in females (J. E. Finnell et al., 2018a).

### 2.5 Drug Information

One hour prior to each WS/CON exposure, females in Experiment #2 received either vehicle (10% dimethyl sulfoxide (DMSO), Fisher Scientific, Pittsburgh, PA) in saline, or the potent and selective ERβ antagonist, 4-[2-Phenyl-5,7-*bis*(trifluoromethyl)pyrazolol[1,5-*a*]pyrimidin-3-yl]phenol (PHTPP, 10 μM dissolved in 10% DMSO, TOCRIS, Minneapolis, MN), microinjected bilaterally into the CeA (0.5 μL over one minute with a one-minute wait post-infusion) (Dennis R. Compton et al., 2004). This dose was chosen based on a dose response pilot in which 10 μM of PHTPP effectively reduced WS-evoked CRF expression in the CeA compared to vehicle and the 1 μM and 3 μM doses of PHTPP (**Figure S1A**).

### 2.6 Behavioral Assays

#### 2.6.1 Behavior During WS/CON

WS/CON sessions were video-recorded on the first (D1) and last (D5) day of exposure, and behaviors including rearing, freezing, and stress-evoked burying (J. E. Finnell et al., 2018a) were manually quantified retrospectively by an experimenter blinded to treatment groups using ANY-maze (Stoelting Co., Wood Dale, IL). Scores were validated by a second blinded experimenter. Briefly, behaviors were defined by the following criteria: stress-evoked burying was frantic, non-exploratory digging or pushing of the bedding; freezing was complete immobility with the exception of respiratory movements and vocalizations; and rearing was upward extension of the rat’s body while simultaneously raising both forepaws from the bedding with or without the hindlimbs extended, in the absence of grooming, eating, or drinking. Importantly, the burying exhibited during stress is a spontaneous and frantic burying of the cage bedding rather than burying of a discrete object. Analyses included the latency to the first occurrence of each behavior as well as the total duration of each behavior throughout the 15-minute session.

#### 2.6.2 Sucrose Preference Test (SPT)

To assess the role for intra-CeA ERβ on WS-induced anhedonia, rats in Experiment #2 underwent a two-bottle choice SPT performed three days pre- as well as three days post-WS/CON exposure. The SPT paradigm utilized in previously published experiments (J. E. Finnell et al., 2018b) was completed with the following modifications: Phase 1: for the first 24 hours, rats were acclimated to the presence of two bottles in their home cage filled with standard drinking water. Phase 2: rats had free access to two standard bottles filled with 1% sucrose in standard drinking water for the next 24 hours. Phase 3: rats were water deprived for 10 hours prior to the start of the dark cycle (1900 hours), when two bottles, one with standard drinking water and one with 1% sucrose, were placed in the cage overnight for 13 hours. The location of the bottles was switched at 2000 hours to rule out the potential for side bias. Total liquid consumption was measured by subtracting bottle weights at the end of the 13 hours from weights at the start of the test. Percent sucrose preference was calculated as follows:

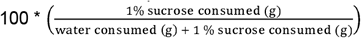

Rats were excluded if they exhibited sucrose aversion indicated by preference below 60% in the pre-stress SPT (CON + VEH, n = 1; CON + PHTPP, n = 2). Data are presented as the change in percent sucrose preference from pre- to post-stress to determine how stress specifically alters this measure of anhedonic behavior.

#### 2.6.3 Acoustic Startle Response (ASR)

ASR testing occurred in a quiet, isolated room using the automated SR-LAB Startle Response System (San Diego Instruments (SDI) San Diego, CA). Briefly, this system consists of a sound-attenuating isolation cabinet which contains an acrylic rat enclosure placed over an accelerometer. To determine how WS specifically alters startle responses, females in Experiment #2 were tested three days prior to and three days following WS/CON exposure. Each session lasted for approximately 30 minutes beginning with an initial 5-minute habituation period of exposure to a constant 65-dB tone (background), followed by 30 total trials that included 10 trials each of 95, 105, and 115 dB tones played in 50-millisecond bursts. Tones were presented in a pseudorandom order with inter-trial intervals that varied between 35-45 seconds. This task elicits an unconditioned startle reflex that has previously been established as a translational method of measuring hypervigilance (Koch & Schnitzler, 1997; Poli & Angrilli, 2015). Since previous experiments indicate that WS impacts startle at the 105 dB level, data were analyzed at 105 dBs as the change in startle velocity (V Max) from pre- to post-WS/CON testing.

#### 2.6.4 Elevated Plus Maze Test (EPM)

Rats in Experiment #2 were tested on the EPM two days following the end of stress. On the day of testing, rats were transported to the testing room at least 30 minutes prior to the start for environmental habituation. The maze was placed 52 cm above the ground and consisted of five total zones: a 10 × 10 cm central square (center zone) that connected two opposing 50 cm x 10 cm extensions enclosed by 40 cm-tall side walls (closed arms) and an additional two opposing 50 × 10 cm extensions without side walls (open arms). Each rat was placed into the center zone facing an open arm, and the location of the animal was tracked for the duration of the 5-minute session through video recording. Videos were automatically scored using ANY-maze to determine the total time spent in each section of the apparatus. This technique provides a measure of anxiety-like/risk averse behavior based on rodents’ inherent aversion to the unfamiliar open spaces (i.e., the open arms) and use of anxiolytics in successfully reducing this aversion (Kraeuter, Guest, & Sarnyai, 2019).

#### 2.6.5 Marble Burying Test (MB)

Rats in both experiments were tested for marble burying, a behavioral task popularly used to assess general anxiety-like behavior in rodents based on defensive burying of a neutral object (Dixit, Sahu, & Mishra, 2020). The testing chamber consisted of a clean, standard rat cage with 5 cm-deep, leveled, Teklad sani-chip bedding – the same as used for all rat’s home cage bedding (Envigo, Dublin, VA). Fifteen glass marbles (1.5 cm in diameter) were arranged in 3 × 5 rows evenly distributed across the bedding (**Figure 3A**). Rats were placed into the cage such that the marbles were not disturbed, and behaviors were video-recorded for 15 minutes. Upon completion, rats were carefully removed so as not to disturb final marble location, and the number of marbles buried was recorded. A marble was counted as buried if ¾ or more of the marble was covered in bedding. Since it is possible that marbles were buried then uncovered, or simply stepped on and appeared buried, durations of marble burying and marble interaction (defined as any physical manipulation of the marble excluding burying) during the 15-minute testing period were manually quantified using ANY-maze. Rats in Experiment #1 underwent a pre- and post-stress test while rats from Experiment #2 underwent only post-stress MB testing. Thus, MB data from Experiment #1 are presented as change from pre-stress while MB data from Experiment #2 represent only behavior during the single MB test conducted following WS exposure.

#### 2.6.6 Witness Stress Context Exposure

On the day of tissue collection, rats from Experiment #1 were exposed to the context in which they originally experienced WS/CON to allow for quantification of stress cue-evoked changes in behavior and neural alterations. Each context exposure was conducted in the same manner as the original WS/CON sessions with the sole exclusion of the resident during WS. Specifically, each female with a history of WS was placed behind a partition in a soiled resident’s home cage with their paired intruder on the opposite side. Females with a history of control-handling were briefly handled then returned to their home cages. Context exposure sessions were video recorded for 15 minutes and behaviors were assessed identically to WS/CON as described above (Section 2.6.1).

### 2.7 Tissue Collection

Brains were collected 30 minutes after the start of WS/CON context exposure for rats in Experiment #1. For rats in Experiment #2, brains were collected at rest two days after the final behavioral test. Rats were deeply anesthetized with isoflurane vapor prior to decapitation. Following dissection, brains were dissected into anterior and posterior sections, flash-frozen in isopentane chilled on dry ice, and stored at -80°C. Brains were sliced coronally, and 1 × 1 mm bilateral punches of the CeA were taken using a tissue biopsy punch (Harvard Apparatus, Holliston, MA) prior to storage at -80°C until processing. Histological assessment verified accuracy of punch placement at 10X magnification (**Figure S1B**).

### 2.8 Brain Tissue Processing

CeA punches were homogenized and assessed for protein concentration using a Pierce Bicinchoninic Acid (BCA) assay (Thermo Scientific, Rockford, IL) per manufacturer’s protocol. Briefly, each sample was mixed with 100 μL of zirconium oxide beads and 200 μL of lab-made lysis buffer consisting of 137 mM NaCl, 20 mM Tris, 1% Ipegal, 10% glycerol, and 1X Halt Protease Inhibitor Cocktail (Thermo Scientific, #87786). Samples then underwent mechanical disruption in a Bullet Blender (Next Advance, Averill Park, NY) for 3 minutes on speed 8 at 4°C followed by centrifugation at 1400 rcf for 15 minutes at 4°C. The resulting homogenate for each sample was mixed in a 1:4 dilution with phosphate buffered sodium azide, and 25 μL of each sample along with 25 μL of each albumin standard was pipetted in duplicate onto a 96-well plate. Working reagent was added at a volume of 200 μL to each well, and the plate was incubated at 37°C for 30 minutes before reading at 562 nm on a Synergy plate reader (Agilent, Santa Clara, CA). Using the resulting protein concentration, homogenates containing 20 μg of protein were aliquoted and stored at -80°C.

### 2.8.1 Western Blotting

Western blots were completed using CeA homogenates in the manner previously described (J. Finnell et al., 2018). Briefly, each sample was mixed at a 1:6 ratio with a solution of β-mercaptoethanol and 6X sample buffer, heated at 75°C for 5 mins, and then loaded into mini-protean gels (Bio-Rad, Hercules, CA). Gels were submerged in 1X running buffer solution (Bio-Rad, Hercules, CA) in electrophoresis chambers for one hour at 135 V. Proteins were then transferred from the gels onto PVDF membranes in 1X transfer buffer (Bio-Rad, Hercules, CA) in transfer chambers for 90 minutes at 100 V. Membranes were then submerged in blocking buffer at room temperature for one hour (1:1 ratio of 1X PBS:Fluorescent Blocking Buffer, MilliporeSigma, Burlington, MA). After blocking, membranes were incubated overnight at 4°C in the following primary antibodies diluted in a solution of blocking buffer with 0.2% Tween-20: mouse anti-GapDH (1:1000, Santa Cruz #sc-365062), rabbit anti-ERβ (1:250, Thermo Fisher #PA1-310B), and rabbit anti-CRF (1:2500, Abcam #ab-184238). The next day, membranes underwent three 10-minute washes in fresh 1X TBS with 10% Tween-20 and were incubated in LI-Cor fluorescent secondary antibodies (IRDye anti-rabbit 800 nm, LI-Cor #926-32213; IR-Dye anti-mouse 680 nm, LI-Cor #962-68072) diluted 1:20,000 in a solution of blocking buffer with 0.2% Tween-20 and 0.1% sodium dodecyl sulfate for one hour. Membranes were washed in TBS with 10% Tween-20 (3 × 10 minutes), imaged on a LI-Cor Odyssey scanner (LI-Cor Biotechnology, Lincoln, NE), quantified using LI-Cor Image Studios Software, and normalized to GapDH protein levels. Importantly, the GapDH protein levels used for normalization did not differ between treatment groups (**Table S1**).

### 2.9 Statistical Analysis

The data included in Experiment #1 were analyzed using unpaired t-tests to compare WS and CON rats, while the dose response data included in the supplement were analyzed via 1-way ANOVA. All behavioral and molecular statistical analyses for Experiment #2 were performed via 2-way ANOVA with Tukey post hoc analyses. Main effects of stress, PHTPP, and stress x PHTPP interactions are provided in the main text results section while post-hoc analyses are indicated by symbols on figures and denoted in figure legends. Outliers were identified and removed if they were more than two standard deviations from the group mean. Data are presented as mean ± standard error of the mean (SEM) with an α = 0.05.

## 3. Results

### 3.1 Experiment #1

#### 3.1.1 Witness Stress Increases Anxiety-Like Behaviors in Females

Experiment #1 determined that females exposed to WS exhibit anxiety-like and hypervigilant behaviors during WS exposure, in post-stress marble burying, and in response to the WS context. WS-exposed rats exhibited higher duration of spontaneous anxiety-like burying of the cage bedding (**Figure 2A**, D1 bury duration: *t*_*10*_ = 2.368, p = 0.0394) and a shorter latency to start burying when compared to CON rats on D1 **(Figure 2B**,, D1 bury latency: *t*_*10*_ = 4.005, p = 0.0025). This trend continued on D5 with increased burying duration (**Figure 2D**, *t*_*10*_ = 3.185, p = 0.0102) and decreased burying latency (**Figure 2E**, *t*_*10*_ = 2.737, p = 0.0210) when comparing WS to CON rats. This behavior was paralleled by a decrease in exploratory rearing behavior in WS rats on D1 (**Figure 2C**, *t*_*10*_ = 3.728, p = 0.0039) but not D5 of stress (**Figure 2F**, *t*_*10*_ = 1.011, p = 0.3357). No differences were observed in freezing duration on either day of stress, with less than five seconds of freezing exhibited by each group (D1: WS 0.84 ± 0.805, CON 2.83 ± 1.52, *t*_*9*_ = 1.165, p = 0.2741; D5: WS 2.8 ± 2.466, CON 3.617 ± 1.837, *t*_*10*_ = 0.2656, p = 0.796, data not shown).

**Figure 2.**
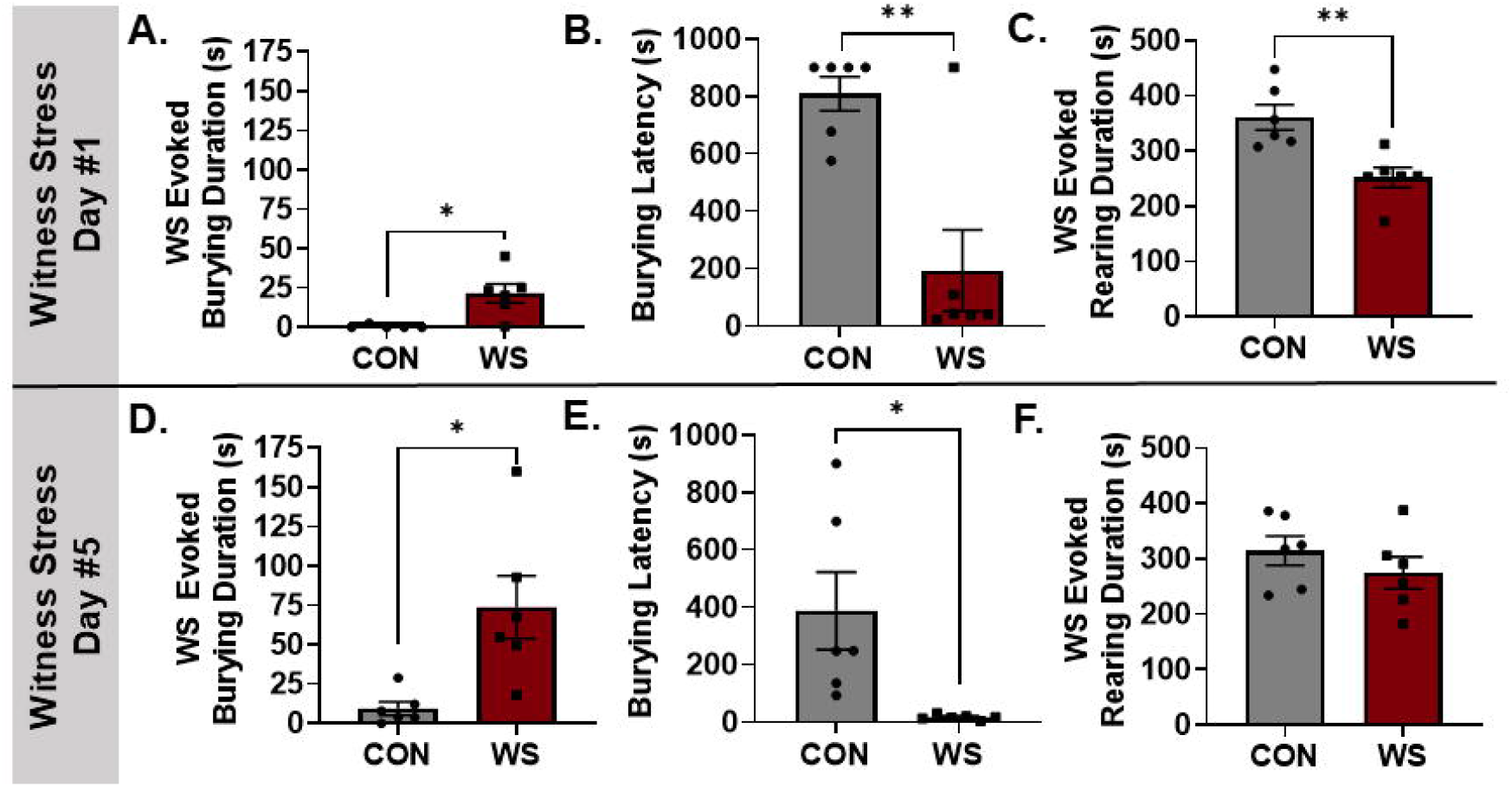
Anxiety-like burying is increased while exploratory rearing is reduced in female rats exposed to WS. On D1, WS rats exhibited an anxiety-like burying response to WS (**A**), a decreased latency to begin burying (**B**), and a reduction in exploratory rearing behavior (**C**). These anxiety-like behavioral effects were also evident following repeated WS exposure as measured on D5, with WS rats exhibiting a heightened burying response (**D**) and decreased burying latency (**E**). On D5, there were no longer reductions in rearing for WS rats compared to CON (**F**) (n = 6/group, \**p* < 0.05, \*\**p* < 0.005). WS: Witness Stress; D1: Day 1; D5: Day 5; CON: control

Following WS/CON exposure, WS rats exclusively exhibited increases in anxiety-like behavior during MB (apparatus depicted in **Figure 3A**). Behaviors including burying, freezing, rearing, and marble interaction were quantified from both the pre- and post-stress tests and compared to achieve a final value reflecting the difference in behavior following stress. Females exposed to WS exhibited an increased number of marbles buried (**Figure 3B**, *t*_*10*_ = 4.385, p = 0.0014) and duration of burying throughout the 15-minute task (**Figure 3C**, *t*_*10*_ = 2.363, p = 0.0397) when compared to CON rats. Additionally, WS-exposed rats exhibited a trend towards decreased time spent interacting with the marbles (**Figure 3D**, *t*_*10*_ = 1.692, p = 0.0608), an exploratory rather than anxiety-like behavior, while CON rats increased their marble interaction. Additional behaviors, including freezing (WS 2.633 ± 2.804, CON 16.33 ± 6.295, *t*_*10*_ = 1.988, p = 0.0749) and rearing (WS 126.9 ± 48.54, CON 36.93 ± 46.33, *t*_*10*_ = 1.34, p = 0.2098), were not different between WS/CON groups (data not shown).

**Figure 3.**
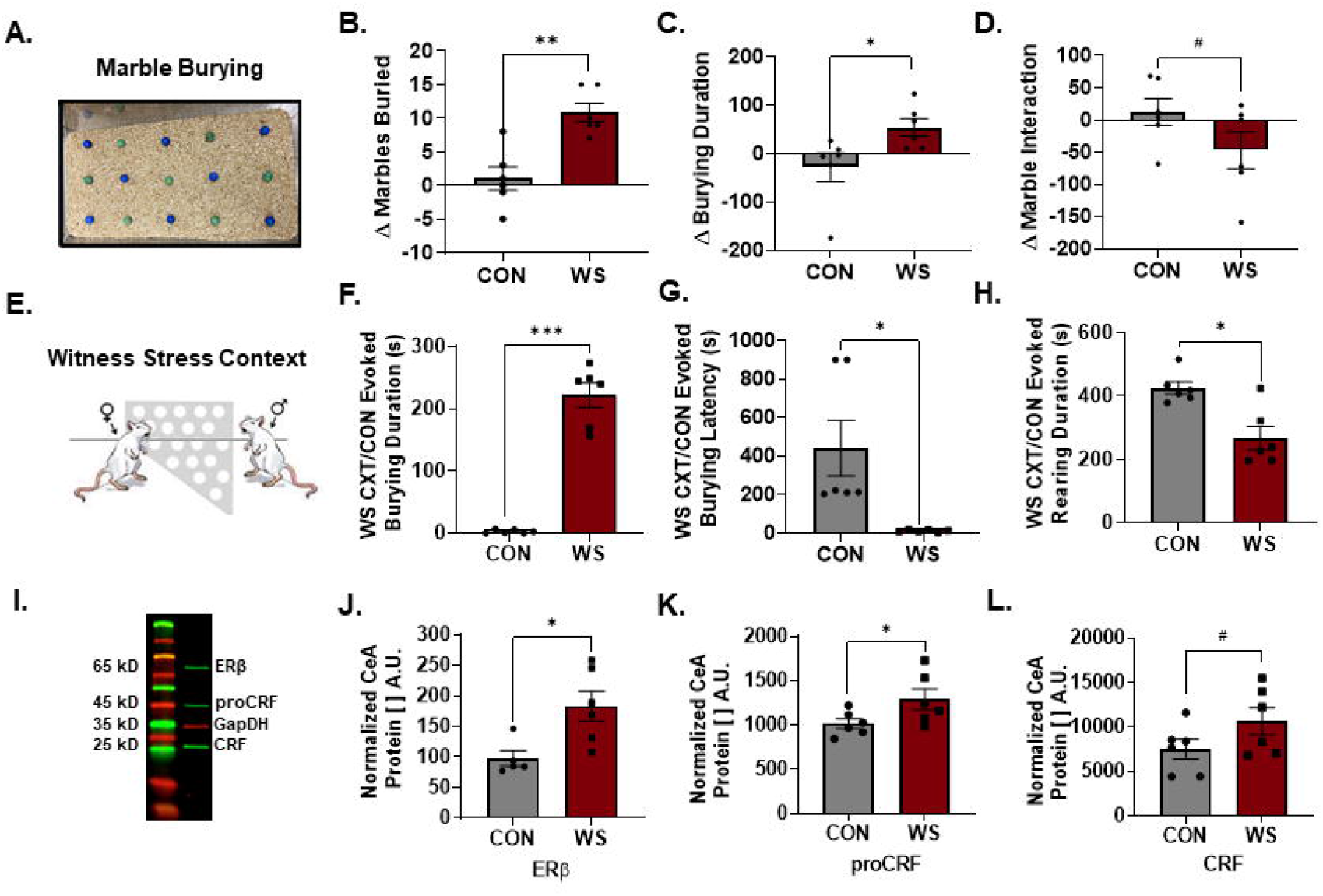
Exposure to social stress increases indices of anxiety-like and hypervigilant behaviors and induces neuronal alterations in the CeA. When compared with pre-stress behaviors in the MB test (apparatus pictured, **A**), WS-exposed rats increased the number of marbles buried (**B**) as well as the overall time spent burying (**C**). This was accompanied by a decrease in the amount of time spent interacting with the marbles (**D**). Further, when exposed to the context and environment in which they originally experienced WS (WS CXT, **E**) versus CON, WS rats exposed to the WS CXT exhibited a significantly heightened amount of time spent spontaneously burying the cage bedding in response to the CXT **(F)** along with a decreased latency to begin burying **(G)** when compared to CON CXT. The enhanced burying response during WS CXT was also accompanied by a reduction in rearing exhibited in WS rats when compared to CON **(H)**. Brain tissue was collected 30 minutes after the start of WS/CON CXT exposure and processed using Western blot analysis for ERβ and CRF normalized to GapDH (**I**). Within the CeA, WS rats displayed augmented expression of ERβ (**J)**, the CRF precursor proCRF (**K**), and the mature CRF protein (**L**) (n = 6/group with the exception statistical of outliers, ^#^*p* < 0.07, \**p* < 0.05, \*\**p* < 0.005, \*\*\**p* < 0.0001, unpaired t-tests). CeA: central amygdala; MB: marble burying; WS: witness stress; CXT: context; CON: control; ERβ: estrogen receptor beta; CRF: corticotropin releasing factor

#### 3.1.2 Exposure to Stress Cues Augments Anxiety-Like Behaviors

The day following MB testing, rats were exposed to their respective WS/CON context (**Figure 3E**) for 15 minutes to mimic their original exposure conditions. When compared to CON rats, which were handled and returned to their home cage for video recording (CON context), rats previously exposed to WS exhibited significantly increased levels of burying (**Figure 3F**, *t*_*10*_ = 11.23, p < 0.0001) along with a decreased latency to start burying (**Figure 3G**, *t*_*10*_ = 2.996, p = 0.0141) in response to the context in which they originally experienced WS. Further, these WS rats exhibited a decrease in rearing behavior (**Figure 3H**, *t*_*10*_ = 3.785, p = 0.0036) when compared to CON.

#### 3.1.3 WS Context Exposure Increases ERβ and CRF in the Central Amygdala

Brain tissue was collected 15 minutes following the end of WS/CON context exposure and brains were prepared for Western blot analysis using antibodies for ERβ and CRF, which yields a separate band for the CRF precursor pro-CRF (**Figure 3I**). WS rats exhibited an increase in ERβ (**Figure 3J**, *t*_*9*_ = 2.928, p = 0.0168) as well as proCRF (**Figure 3K**, *t*_*10*_ = 2.113, p = 0.0304) with a trend towards increased CRF (**Figure 3L**, *t*_*10*_ = 1.648, p = 0.0652) in the CeA in response to the WS context.

### 3.2 Experiment #2

#### 3.2.1 WS-Evoked Burying is Attenuated by Intra-CeA ERβ Antagonism during Repeated WS Exposure

Similar to previous findings in our lab and in accordance with Experiment #1, WS was shown to promote anxiety-like burying on D1 (**Figure 4A**, main effect of stress: *F*_*1,41*_ = 32.83, *p* < 0.0001) and D5 of WS/CON exposure (**Figure 4D**, main effect of stress, *F*_*1,32*_ = 19.17, *p* =0.0001) along with a stress x drug interaction observed on D5 (interaction, *F*_*1,32*_ = 4.209, *p* =0.0485). Specifically, while PHTPP had no effect on the burying response upon the initial WS exposure (D1 *post hoc*, WS + VEH vs. WS + PHTPP, p = 0.6135), ERβ blockade significantly reduced WS-evoked burying on D5 following repeated WS exposure (Day #5 *post hoc*, WS + VEH vs. WS + PHTPP, p = 0.0239). Further, WS rats exhibited a shorter latency to begin burying on both D1 (**Figure 4B**, main effect of stress: *F*_*1,45*_ = 38.49, *p* < 0.0001) and D5 (**Figure 4E**, main effect of stress: *F*_*1,36*_ = 68.27, *p* < 0.0001). Additionally, there were no significant main effects observed regarding rearing on D1 (**Figure 4C**, stress effect, *F*_*1,46*_ = 0.0067, *p* = 0.9351, drug effect, *F*_*1,46*_ = 0.5575, *p* = 0.4591) or D5 (**Figure 4F**, stress effect, *F*_*1,37*_ = 0.0103, *p* = 0.9197, drug effect, *F*_*1,37*_ = 1.791, *p* = 0.1890) as well as freezing on D1 (stress effect, *F*_*1,46*_ = 1.957, *p* = 0.1685, drug effect, *F*_*1,46*_ = 0.07395, *p* = 0.7869) or D5 (stress effect, *F*_*1,34*_ = 0.1455, *p* = 0.7052, drug effect, *F*_*1,34*_ = 0.00281, *p* = 0.958, data not shown).

**Figure 4.**
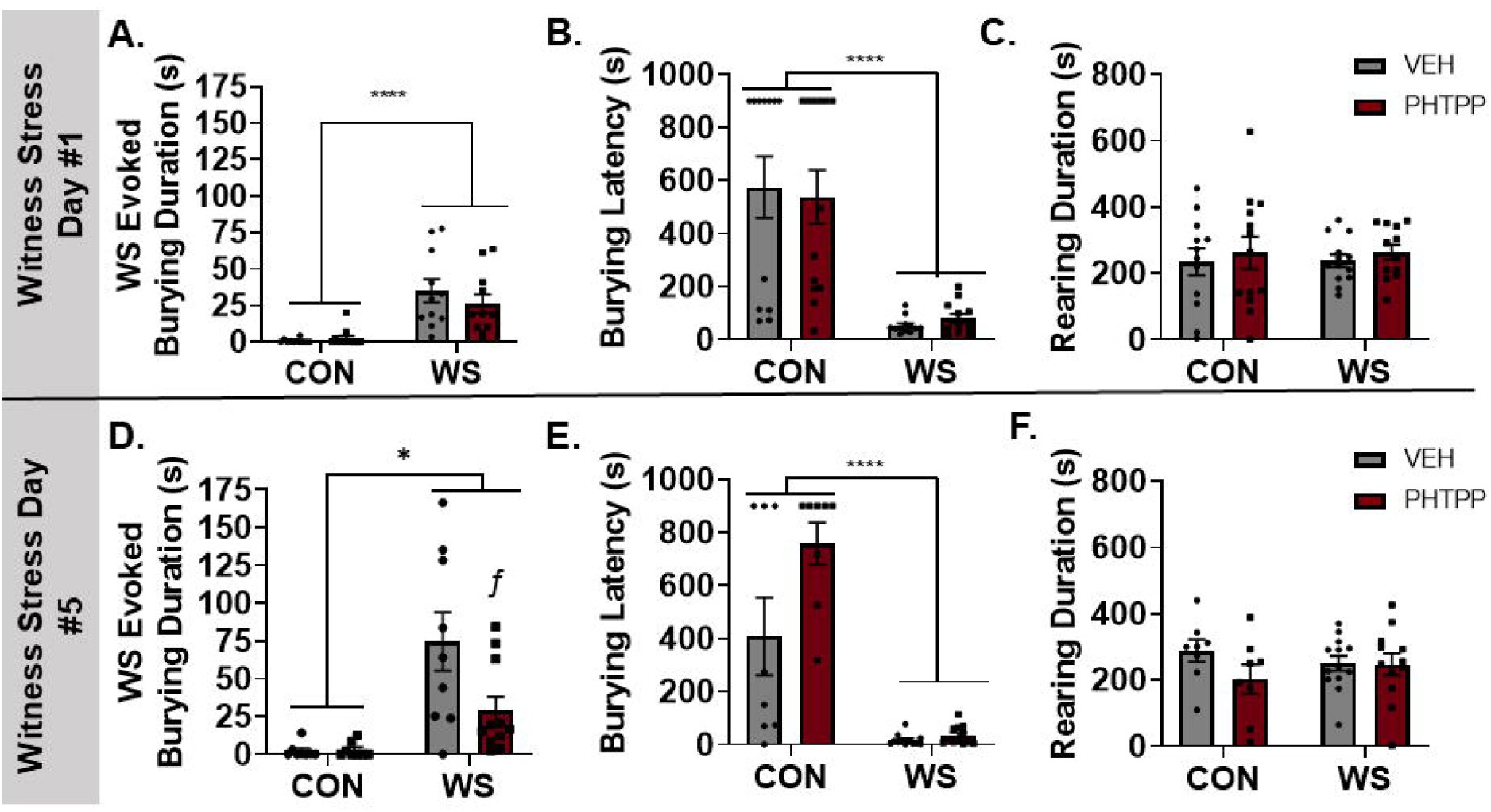
Blocking ERβ in the CeA reduces spontaneous WS-evoked burying during repeated WS exposure. Spontaneous burying behavior was increased among WS rats compared to CON during day #1 (D1) exposure to WS/CON, and this acute response was unaffected by pre-treatment with the ERβ antagonist PHTPP (**A**). Burying also presented significantly quicker in WS rats (**B**) without any group differences in rearing (**C**). However, on the fifth day of WS exposure, PHTPP attenuates the WS-evoked burying response (**D**, ^−^ *p* < 0.05, WS + VEH vs. WS + PHTPP) along with an overall stress effect on latency to begin burying and no significant differences between the groups with regards to rearing behavior (n = 12-13/group with the exception of statistical outliers, \**p* < 0.05, \*\*\*\**p* < 0.0001). ERβ: estrogen receptor beta; CeA: central amygdala; WS: witness stress; D1: day 1; CON: control

#### 3.2.2 WS and PHTPP Differentially Regulate Anxiety-like Behaviors

Following WS/CON, rats were tested on a variety of behavioral paradigms to determine the impact of WS and/or ERβ antagonism on the behavioral adaptations that occur in response to repeated stress. In the EPM, repeated WS exposure resulted in a reduction in the percentage of time spent in the open arms (**Figure 5A**; main effect of stress: *F*_*1,37*_ = 4.231, *p* = 0.0468). Importantly, there was no overall effect on the total distance traveled (**Figure 5B**, stress effect, *F*_*1,35*_ = 0.0379, *p* = 0.8468, drug effect, *F*_*1,37*_ = 0.1983, *p* = 0.6588). However, while there was a significant stress effect on this measure of anxiety-like behavior, there was no effect of PHTPP (main effect of drug: *F*_*1,37*_ = 0.1515, *p* = 0.6993). Additionally, the percentage of time spent in the closed arms or center zone was not affected by stress (closed arms: *F*_*1,35*_ = 1.228, *p* = 0.2753;, center: *F*_*1,35*_ = 1.087, *p* = 0.3044) nor PHTPP (closed arms: *F*_*1,35*_ = 0.3156, *p* = 0.5778; center: *F*_*1,35*_ = 0.4888, *p* = 0.4891, data not shown). When tested for ASR, repeated WS exposure resulted in an enhanced startle amplitude, specifically at 105 dB, and this exaggerated startle response is diminished in WS rats that were treated with PHTPP (**Figure 5C**; stress x drug interaction: *F*_*1,24*_ = 17.13, *p* = 0.0004). Specifically, amongst vehicle-treated rats, there was a significant increase in startle amplitude in the WS rats (CON + VEH vs. WS + VEH, p = 0.0015), and this was significantly blunted by PHTPP treatment during WS (WS + VEH vs. WS + PHTPP, p = 0.0002). In the subsequent testing to determine negative valence-related behaviors, there was a significant stress x drug interaction (**Figure 5D**, *F*_*1,30*_ = 4.674, *p* = 0.0387) by which sucrose preference was significantly reduced following WS exposure, though PHTPP treatment during WS exposure prevented this reduction (WS + VEH vs. WS + PHTPP, p = 0.0278). Finally, MB revealed a trend toward a stress x drug interaction (*F*_*1,32*_ = 3.284, *p* = 0.079) with a significant drug effect (*F*_*1,32*_ = 8.679, *p* = 0.006) where PHTPP-treated WS rats exhibited a decrease in the number of marbles buried compared to vehicle-treated WS rats (**Figure 5E**, *p* = 0.016). This effect was also observed in the overall time spent burying during MB, with a significant drug effect (*F*_*1,28*_ = 7.667, *p* = 0.0099), trend toward a stress x drug interaction (*F*_*1,28*_ = 3.974, *p* = 0.056), and significant difference between WS + VEH- and WS + PHTPP-treated rats (**Figure 5F**, *p* = 0.0182). There were no significant effects of stress or drug on additional behaviors recorded during this task including marble interaction (stress effect: *F*_*1,31*_ = 0.02224, *p* = 0.8819; drug effect: *F*_*1,31*_ = 1.567, *p* = 0.22, data not shown). It is important to note that these data represent behaviors from a single post-stress task which may explain their difference from the robust results seen in the pre- to post-stress comparisons in Experiment #1.

**Figure 5.**
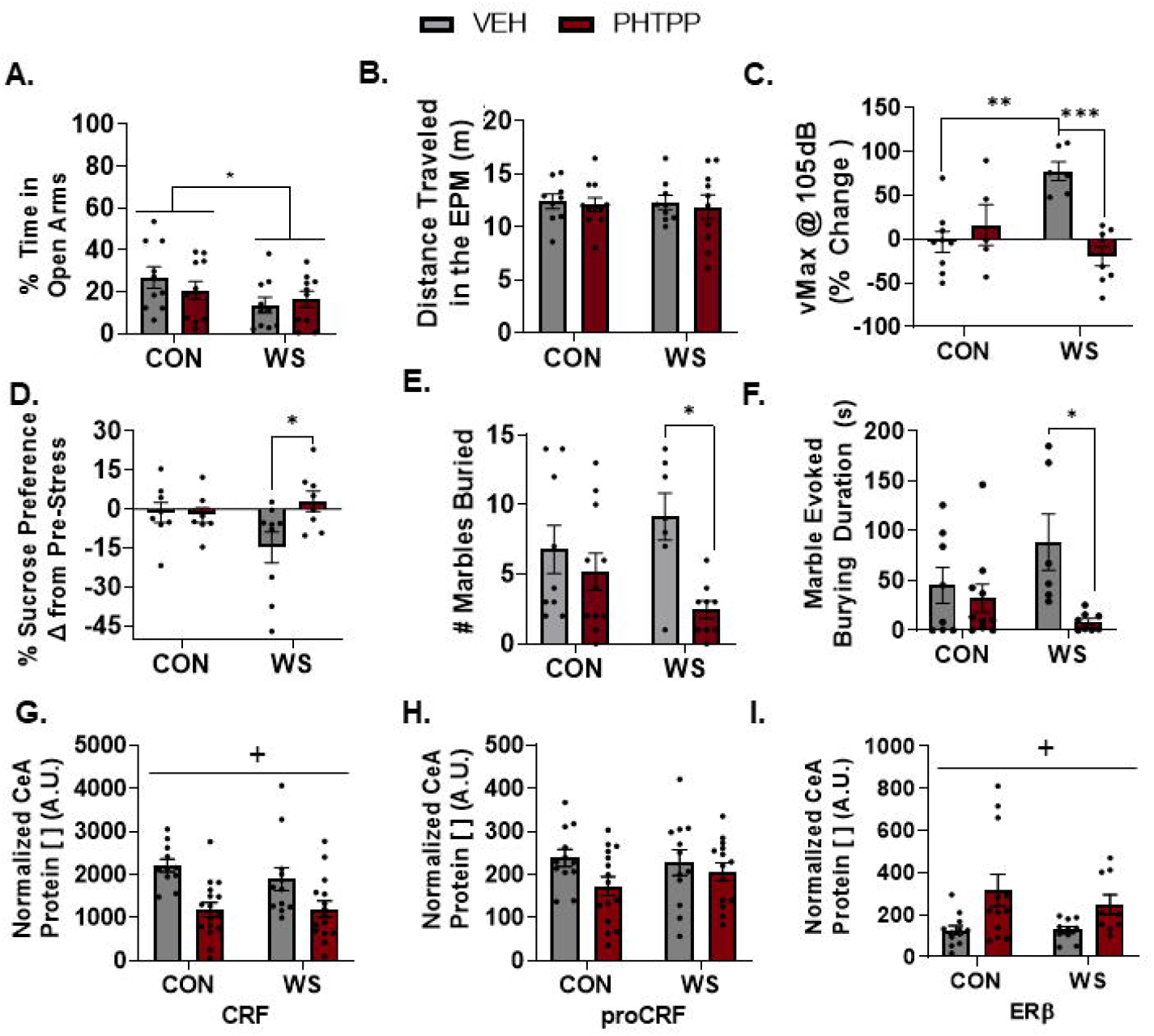
A history of WS and ERβ antagonism in the CeA differentially affect hypervigilant, anxiety- and depressive-like behaviors. Repeated WS produces persistent increases in anxiety-like behavior evidenced by a modest reduction in % time in the open arm during EPM (**A**) with no difference in motor behaviors evidenced by overall distance traveled (**B**). Modest anxiety-like behavior displayed on the EPM was not regulated by pharmacological blockade of ERβ during WS exposure, as evidenced by a lack of effect induced by PHTPP. Further, a history of repeated WS rendered rats with a hypervigilant phenotype indicated by an enhanced acoustic startle response. Post-WS startle was increased by WS rats compared to CON and blocking ERβ during the WS episodes with PHTPP prevented the development of heightened ASR (**C**). A history of repeated WS exposure resulted in a significant decrease in the percent of sucrose preferred in the post-stress test but this reduction in reward seeking/depressive-like behaviors was prevented by PHTPP (**D**). Behaviors measured in the final task, MB, revealed a significant effect of PHTPP treatment, with WS rats treated with PHTPP exhibiting a significant decrease in the number of marbles buried (**E**) as well as in the overall time spent burying the marbles (**F**) when compared to vehicle treated WS rats. Brains were collected under resting conditions 48 hours after the final marble burying task. CeA tissue was prepared for Western blot analysis; PHTPP caused a compensatory increase in ERβ at rest in both control and WS rats (**G**), with no effect on proCRF (**H**), and a significant long-term reduction in resting levels of CRF (**I**) (n = 12-13/group with the exception of statistical outliers, \**p* < 0.05, \*\**p* < 0.005, \*\*\**p* < 0.0005, \*\*\*\**p* < 0.0001). WS: witness stress; ERβ: estrogen receptor beta; CeA: central amygdala; EPM: elevated plus maze; ASR: acoustic startle response; MB: marble burying; CRF: corticotropin releasing factor

#### 3.2.3 PHTPP Induces Long-Term Alterations in CeA CRF and ERβ

Tissue from Experiment #2 was collected under resting conditions two days following the final behavioral assay. Brain samples were analyzed using Western blot to establish protein expression for ERβ and CRF. There was a significant drug effect of previous PHTPP treatment on long-term resting ERβ protein expression in the CeA (**Figure 5G**, *F*_*1,39*_ = 10.11, *p* = 0.0029) with PHTPP-treated rats exhibiting higher levels of ERβ when compared to vehicle, independent of stress history. Additionally, PHTPP treatment led to decreases in CRF protein expression in the CeA for both WS and CON groups (**Figure 5I**, main effect of drug, *F*_*1,39*_ = 18.76, *p* < 0.0001) with no significant change in proCRF (**Figure 5H**, stress x drug interaction, *F*_*1,49*_ = 0.8941, *p* = 0.349).

## 4. Discussion

These experiments were completed to determine the contribution of CeA ERβ signaling on stress-induced behavioral outcomes due to the prominent role of the CeA in stress responsivity and the dichotomous role of estrogen on behavior. Overall, these data show that WS increases negative valence behaviors that were accompanied by increases in intra-CeA ERβ and CRF protein expression, though pharmacological blockade of ERβ in the CeA prevented these effects. These studies build on previous work investigating the role of central estrogen signaling on stress-related outcomes in females and highlight a novel treatment target for stress-induced neuropsychiatric disorders.

The increased prevalence of stress-related psychiatric disorders among women in their reproductive years is well documented (Barth, Villringer, & Sacher, 2015; Bezerra, Alves, Nunes, & Barbosa, 2021) and, while we cannot rule out contributing roles of other ovarian hormones such as progesterone, accumulating evidence from both clinical (Hlavacova, Wawruch, Tisonova, & Jezova, 2008; Newhouse et al., 2010) and preclinical (Figueiredo, Ulrich-Lai, Choi, & Herman, 2007; Flores et al., 2020; Hokenson et al., 2021; Kokane & Perrotti, 2020; Shansky et al., 2004) studies demonstrate that, in the context of stress, central estrogen signaling plays a critical role in this heightened susceptibility. It is reasonable, then, to consider estrogen signaling through ERβ within the CeA resulting in increased CRF expression as a promising mechanism by which estrogen facilitates this enhanced stress response among females. Our findings support this hypothesis as the first to identify a regulatory role for intra-CeA ERβ in the development of negative valence phenotypes that emerge among naturally cycling females as a consequence of repeated social stress exposure.

While ERβ blockade within the CeA during WS exposure protected against behavioral sensitization to WS, it is important to consider the impact of this pharmacological manipulation on CeA projection regions that are known to directly regulate these behaviors. Importantly, CRF-projecting neurons from the CeA have been shown to activate noradrenergic neurons of the LC (Curtis, Bello, Connolly, & Valentino, 2002; Curtis, Lechner, Pavcovich, & Valentino, 1997), which is responsible for initiating the hypervigilant burying behavior assessed in our studies (Howard et al., 2008). Given the role for LC-norepinephrine (NE) signaling in CRF-induced burying responses, reduced LC-NE signaling due to decreased stress-induced CRF expression in the PHTPP-treated rats could explain their blunted WS-evoked burying response (J. E. Finnell et al., 2018a). Unfortunately, the time point of euthanasia in these experiments does not allow for the analysis of this hypothesis within our current dataset. Future studies exploring this avenue include dynamic methods of analysis, like *in vivo* electrophysiology, to better understand the impact of CeA ERβ antagonism on LC-NE signaling and behavior.

It is also important to note that while intra-CeA ERβ appears to mediate many stress-induced behaviors, EPM was unaffected by intra-CeA PHTPP treatment. While WS rats regardless of drug treatment exhibited a more risk-averse phenotype in this task, ERβ blockade during WS failed to elicit any effects. This may be due to the involvement of other brain regions in the regulation of EPM-specific behaviors, including the medial amygdala (Troakes & Ingram, 2009) and bed nucleus of the stria terminalis (Butler et al., 2016; Callahan, Tschetter, & Ronan, 2013), and, thus, sole modulation of ERβ in the CeA is insufficient in altering these behaviors. An additional unexpected finding was that, in the long-term, treatment with PHTPP for five consecutive days resulted in a compensatory resting increase in ERβ expression within the CeA. While PHTPP is established as a potent and selective antagonist of ERβ (D. R. Compton et al., 2004), there is a lack of research examining neuronal adaptations that occur in response to repeated pharmacological blockade of ERβ. While these are the first results to show that five days of PHTPP microinjection into the CeA lead to compensatory increases in receptor expression, further studies should detail the time course of this alteration as well as any additional latent behavioral effects.

Overall, these experiments have established that WS exposure increases intra-CeA CRF expression and promotes the development of negative valence behaviors across multiple behavioral paradigms. Further, despite the compensatory elevation in CeA ERβ, local PHTPP treatment during WS exposure was effective in attenuating the stress-induced neuronal and behavioral shifts identified in these experiments. Future studies that examine downstream neural correlates involved in these behavioral responses and impacted by ERβ signaling, such as the LC-NE system, that may serve as viable targets for the treatment and prevention of stress-induced neuropsychiatric disorders in females.

## Conclusions

These experiments are the first to demonstrate that repeated exposure to witness stress results in an increase of ERβ and CRF expression within the CeA, and further, that pharmacological blockade of intra-CeA ERβ during each stress exposure prevents the emergence of behavioral deficits indicative of negative valence phenotypes as measured by marble burying, acoustic startle, and sucrose preference.

## Abbreviations

CeA: central amygdala
CRF: corticotropin releasing factor
WS: witness stress
ERβ: estrogen receptor beta
PHTPP: 4-[2-Phenyl-5, 7-bis (trifluoromethyl) pyrazolo [1,5-a] pyrimidine-3-yl] phenol
CORT: corticosterone
HPA: hypothalamic pituitary adrenal axis
ACTH: adrenocorticotropic hormone
CON: control handing
SPT: sucrose preference testing
ASR: acoustic startle responding
EPM: elevated plus maze
dB: decibels
MB: marble burying
BCA: bicinchoninic acid
LC: locus coeruleus

