## Supplemental Information for "Estrogen receptor beta in the central amygdala regulates the deleterious behavioral and neuronal consequences of repeated social stress in female rats"

**PHTPP Dose Response Curve:** An initial pilot experiment was completed to determine the optimal concentration of the estrogen receptor beta (ERβ) antagonist, PHTPP, for reducing witness stress (WS)-evoked corticotropin releasing factor (CRF) expression within the central amygdala (CeA). Rats were implanted with indwelling bilateral microinjection cannulas aimed at the CeA and given at least 7 days for surgical recovery. PHTPP was prepared at concentrations of 1, 3, or 10 μM in 10% dimethyl sulfoxide (DMSO) and rats were given CeA microinjections of vehicle (10% DMSO) or PHTPP one hour prior to a single WS exposure (15 minutes). Animals were sacrificed 90 minutes following the start of WS and brains were prepared for CRF analysis via Western Blot.

**Supplemental Figure 1. A)** PHTPP attenuates WS-evoked increases in CeA CRF levels at a concentration of 10 μM while the 1 and 3 μM injections were insufficient. Thus, all experiments utilizing PHTPP were completed using a concentration of 10 μM. **B)** For all experiments, brains were processed for collection of the CeA (detailed methods in main text). All punches were assessed to maintain anatomical localization within the CeA, and a representative image of correct placement is presented here. **C)** The hypothesized mechanism through which estrogen increases stress susceptibility in females that experiments within the manuscript were based on**.** Receptor bound estrogen in the CeA has the potential to bind to an estrogen response element on the CRF gene which promotes the transcription of CRF mRNA. This increase in CRF as a result of estrogen binding to ERβ results in an augmented HPA axis response and increased stress responsivity/sensitization.

**Supplemental Figure 2. A)** Western blot images for the data in Figure 3.  **B)** Western blot images for the data in Figure 5.

**Supplemental Table 1.** Verification of GapDH protein levels by treatment group for all analyses.
