## Supplementary figures and images for "Estrogen receptor beta in the central amygdala regulates the deleterious behavioral and neuronal consequences of repeated social stress in female rats"

### Supplemental Figure 1

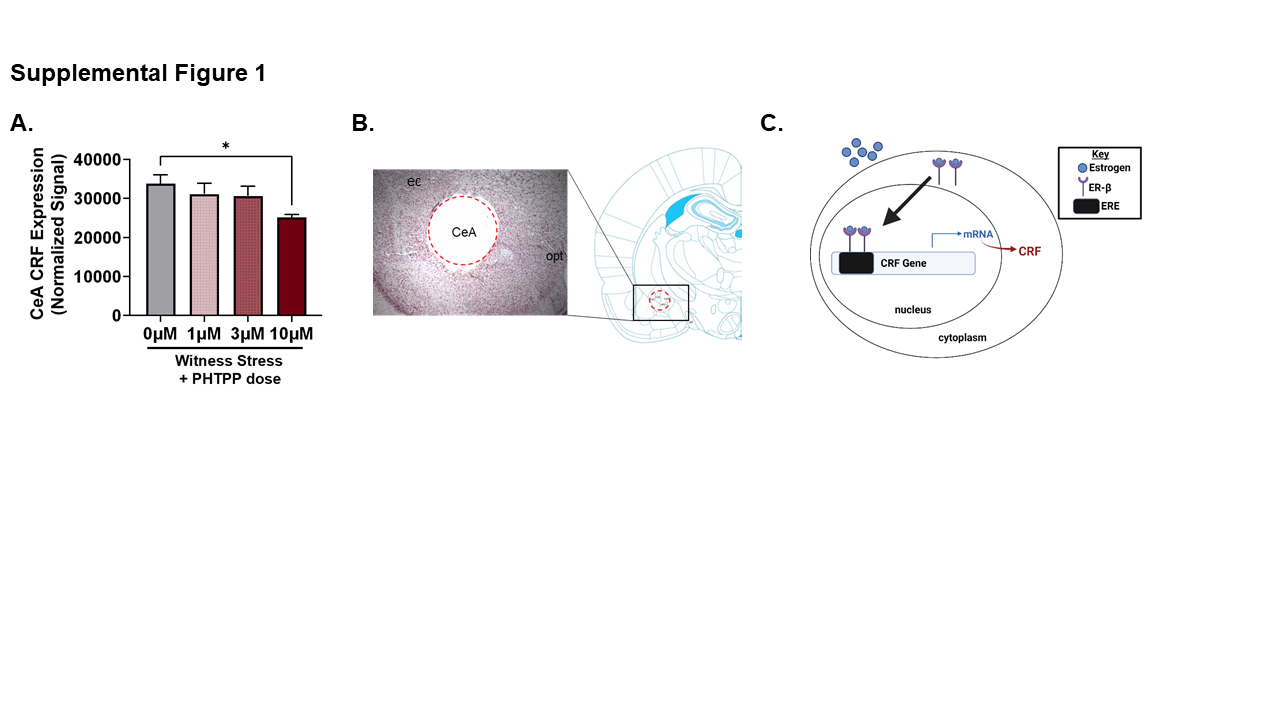

### Supplemental Figure 2

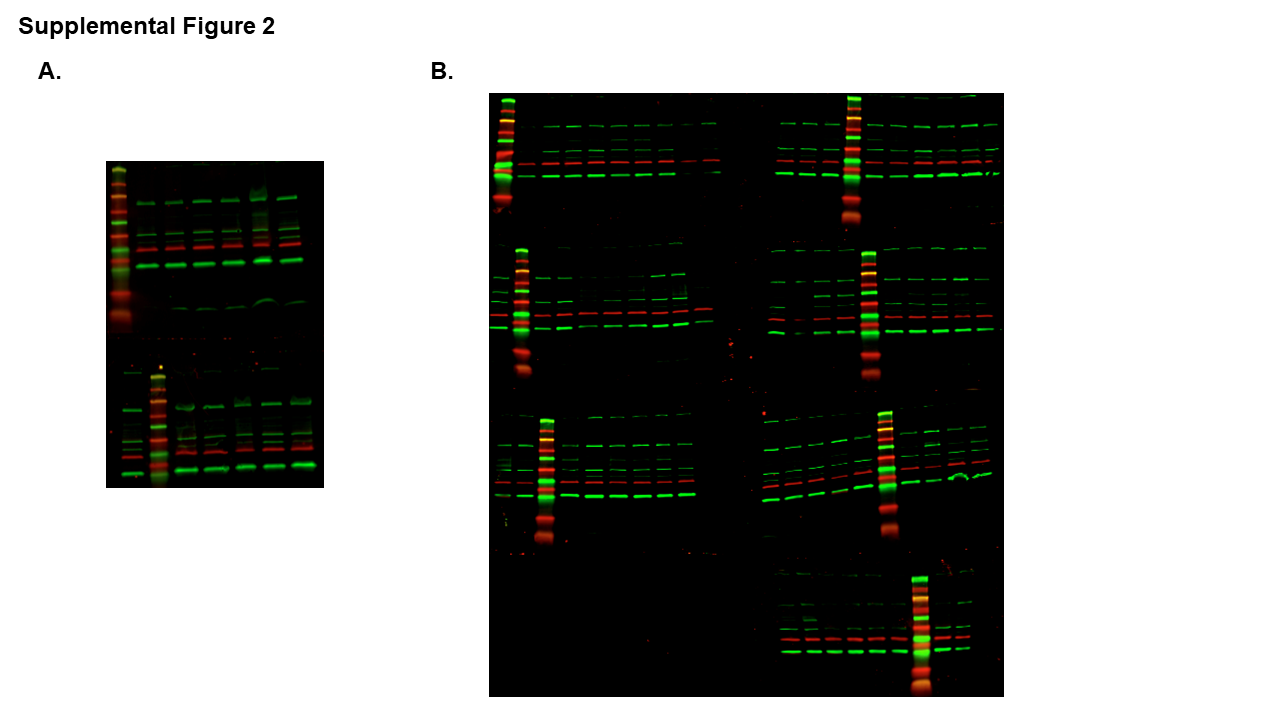

### Supplemental Table 1

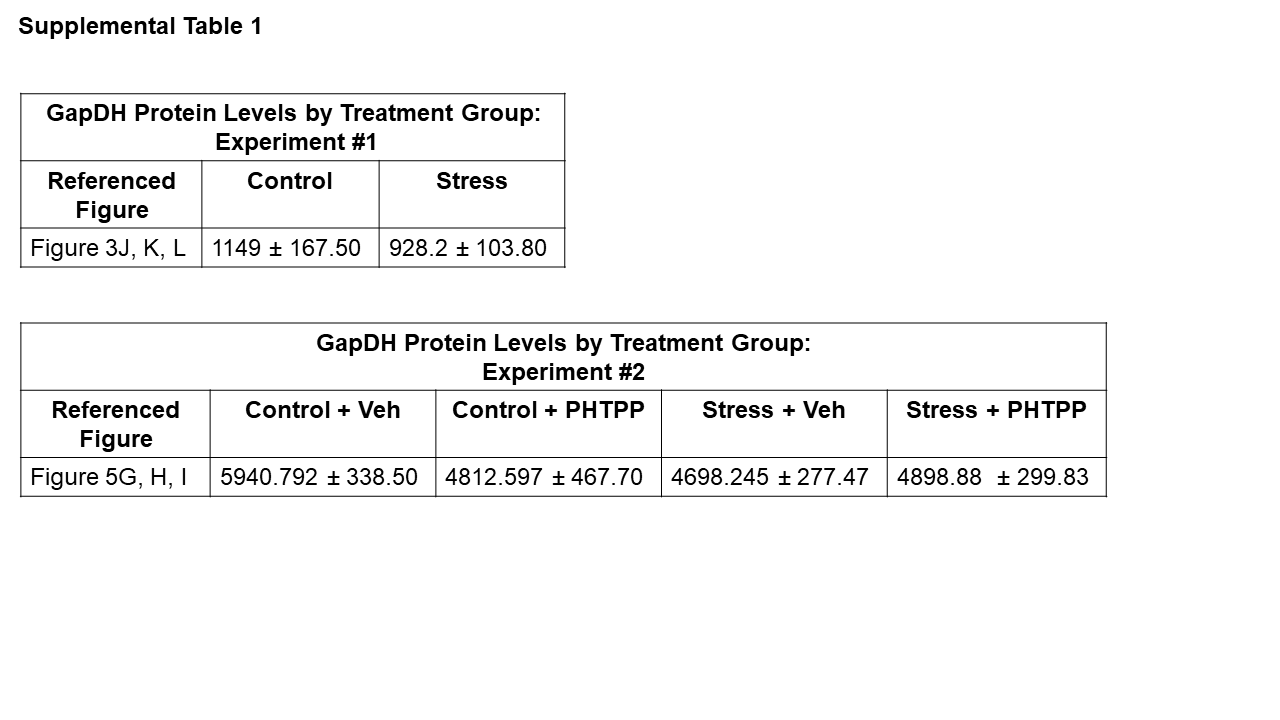
